# EXORIBONUCLEASE4 integrates metabolic signals induced by osmotic stress into the circadian system

**DOI:** 10.1101/2022.07.05.498805

**Authors:** Putri Prasetyaningrum, Suzanne Litthauer, Franco Vegliani, Matthew William Wood, Martin William Battle, Cathryn Dickson, Matthew Alan Jones

## Abstract

The circadian clock system acts as an endogenous timing reference that coordinates many metabolic and physiological processes in plants. Previous studies have shown that the application of osmotic stress delays circadian rhythms via 3’-Phospho-Adenosine 5’-Phosphate (PAP), a retrograde signalling metabolite that is produced in response to redox stress within organelles. PAP accumulation leads to the inhibition of EXORIBONUCLEASEs (XRNs), which are responsible for RNA degradation. Interestingly, we are now able to demonstrate that post-transcriptional processing is crucial for the circadian response to osmotic stress. Our data show that degradation of specific circadian clock transcripts is modulated by osmotic stress, suggesting that RNA metabolism plays a vital role in circadian clock coordination during drought. Inactivation of XRN4 is sufficient to extend circadian rhythms, with *LWD1, LWD2*, and *PRR7* identified as specific XRN4 targets that are post-transcriptionally regulated to delay circadian progression.

**One Sentence Summary:** Post-transcriptional regulation of specific transcripts enables the circadian system to respond to osmotic stress.

## Introduction

Drought is one of the primary contributors to the yield gap that exists between theoretical yields and those realised in the field (Gupta et al., 2020). However, plants’ responses to drought are complex and so our understanding of underlying signaling pathways remains limited. One of the initial steps in plants’ perception of drought stress is the induction of oxidative stress in the chloroplast (Chan et al., 2016a). These stresses lead to the inactivation of the redox-sensitive enzyme SAL1, resulting in the accumulation of 3’-PhosphoAdenosine 5’-Phosphate (PAP), a retrograde signalling molecule that indicates metabolic stress within the chloroplast (Chan et al., 2016a; Koprivova and Kopriva, 2016; Phua et al., 2018). The accumulation of PAP alters global patterns of transcription and RNA catabolism by inhibiting the activity of 5’-3’ exoribonucleases [XRNs; (Gy et al., 2007; Kurihara et al., 2012; Crisp et al., 2018)], while PAP also serves as a secondary messenger to promote abscisic acid (ABA) signalling (Pornsiriwong et al., 2017). Higher order Arabidopsis mutants lacking all three Arabidopsis XRNs (XRN2, XRN3, XRN4; *xrn234*) have improved drought tolerance, similar to mutant lines that constitutively accumulate PAP (Hirsch et al., 2011). However, these higher order *xrn234* mutants are unlikely candidates for crop improvement as such plants grow slowly and have a delayed flowering phenotype (Hirsch et al., 2011). By comparison, loss of *XRN4* has modest effects on plant growth and physiology, although roles in seed germination and impaired responses to hormones including ethylene, ABA, and auxin have been reported (Basbouss-Serhal et al., 2017; Wawer et al., 2018; Windels and Bucher, 2018). Instead, XRN4 appears to have a greater role in mediating plants’ responses to abiotic factors including heat and salt stress by contributing to cytosolic RNA degradation (Merret et al., 2013; Merret et al., 2015; Nguyen et al., 2015; Kawa et al., 2020).

Mature mRNAs are protected from degradation by a 5’ 7-methylguanosine cap and the 3’ polyadenosine tail, with polyadenylation serving as the primary determinant of degradation rate (Decker and Parker, 1993; Sieburth and Vincent, 2018). Beyond these initial regulatory steps, cytosolic RNA degradation can occur in either a 3’ →5’ or 5’→3’ direction. The exosome and SUPRESSOR OF VARICOSE (SOV) contribute to 3’→5’ degradation, whereas XRN4 is the primary actor in 5’→3’ degradation; XRN2 and XRN3 are exclusively localized to the nucleus (Nagarajan et al., 2013; Sieburth and Vincent, 2018). These pathways are broadly conserved across metazoans and fungi, although animals and fungi utilize a functionally equivalent XRN4 orthologue (XRN1) for cytosolic 5’→3’ degradation (Kastenmayer and Green, 2000; Nagarajan et al., 2013). Degradation of cytoplasmic RNA via these pathways prevent the generation of siRNAs (Sieburth and Vincent, 2018; Zhang et al., 2015), although the function of these partially-degraded intermediates remain otherwise unstudied. Instead, recent reports demonstrate unanticipated relationships between these conserved RNA degradation pathways and other aspects of RNA metabolism and processing in yeast (Blasco-Moreno *et al.*, 2019; Haimovich *et al.*, 2013; Sun *et al.*, 2013). In plants, at least a portion of XRN4-mediated degradation occurs co-translationally, leading to *xrn4* seedlings having impaired translation of specific transcripts (Carpentier *et al.*, 2020).

We have previously reported that the accumulation of PAP leads to a delay in the circadian system, and that comparable phenotypes are observed in *xrn234* seedlings (Litthauer et al., 2018). The circadian system is a molecular timekeeping mechanism that enables time-of-day to be integrated into plants’ responses to environmental signals (Millar, 2016). Timing information provided by the circadian system allows anticipation of regular environmental changes (such as dawn and dusk) whilst also modulating gene expression in response to stresses (Greenham and McClung, 2015; Grundy et al., 2015; Baek et al., 2020). Manipulation of the circadian system can improve drought tolerance (Legnaioli *et al.*, 2009, Nakamichi *et al.*, 2016) and alter water use efficiency (Simon et al., 2020). Indeed, the circadian system has been proposed as a key target to improve agronomic traits in breeding programs (Bendix et al., 2015). We therefore sought to determine how osmotic stress contributes to the regulation of circadian timing.

The circadian system is multi-faceted but relies on interlocking transcriptional negative feedback loops that generate daily rhythms of approximately 24 hours (Millar, 2016). Morning-phased components, such as CIRCADIAN CLOCK ASSOCIATED1 (CCA1) work in combination with PSEUDO RESPONSE REGULATOR9 (PRR9), PRR7, and PRR5 to repress gene expression throughout the day (Alabadi et al., 2001; Nakamichi et al., 2010). At night, the Evening Complex [primarily comprising of EARLY FLOWERING3 (ELF3), ELF4, and LUX ARRHYTHMO (LUX)] inhibits gene expression (Nusinow et al., 2011; Huang et al., 2016). These waves of repression are complemented by transcriptional activators including LIGHT-REGULATED WD1 (LWD1) and LWD2 that promote expression of morning-phased clock genes (Wang et al., 2011; Wu et al., 2016). Following transcription, proteins such as GIGANTEA contribute to the post-translational regulation of circadian timing (Kim et al., 2007; Cha et al., 2017)

Although primarily examined at the transcriptional level, the contribution of post-transcriptional regulation to the maintenance of circadian rhythms is becoming apparent. Alternative splicing, nuclear export, and Non-sense Mediated Decay (NMD) all contribute to circadian timing, and *CCA1* transcript has been reported to be less stable in the presence of light (Yakir et al. 2007; Jones et al., 2012; Wang et al., 2012; Macgregor et al., 2013; Kwon et al., 2014; Nolte and Staiger, 2015; Romanowski and Yanovsky, 2015; Mateos et al., 2018). Equally, post-transcriptional regulation is similarly recognized as contributing to plants’ responses to abiotic stress (Filichkin et al., 2015; James et al., 2018). In this study, we demonstrate that loss of XRN4 activity is sufficient to delay circadian timing, and that XRN4 contributes to the degradation of *PRR7* and *LWD1.* Importantly, neither *prr7* or *lwd1lwd2* seedlings are able to delay their circadian system in response to osmotic stress, demonstrating how signals from environmental stresses can be integrated into the circadian system.

## Results

### Circadian responses to osmotic stress occur at transcriptional and post-transcriptional levels

We were interested how different aspects of the circadian system responded to osmotic stress and so we utilised a catalogue of luciferase reporter lines to monitor circadian rhythms following transfer to 200mM mannitol (Figure 1, Supplemental Figure 1). Osmotic stress significantly extends circadian free-running period (FRP) under constant red and blue light using luciferase reporters driven by the promoter of *CCA1* or *GI,* in line with our initial studies [Figure 1A-C, (Litthauer et al., 2018)]. However, we were interested to note that *pPRR9::LUC2* and *pLWD1::LUC2* reporter lines did not demonstrate an extension of FRP following osmotic stress (p=0.79 and p=0.50 respectively, Figures 1D-E). Indeed, although circadian parameters were able to be determined for *pLWD1::LUC2* and *pLWD2::LUC2* after the transfer to 200 mM mannitol these oscillations were greatly damped (and presented an increased Relative Amplitude Error; RAE), suggesting that expression of these genes is driven to functional arrhythmia in the presence of osmotic stress (Figures 1E-1F, Supplemental Figure 1G and 1I). *pLWD2::LUC2* lines retained an extension of circadian period following the application of osmotic stress (Figure 1F). Such data demonstrate that promoter activity of a subset of circadian genes is disrupted following the application of osmotic stress.

**Figure 1.**
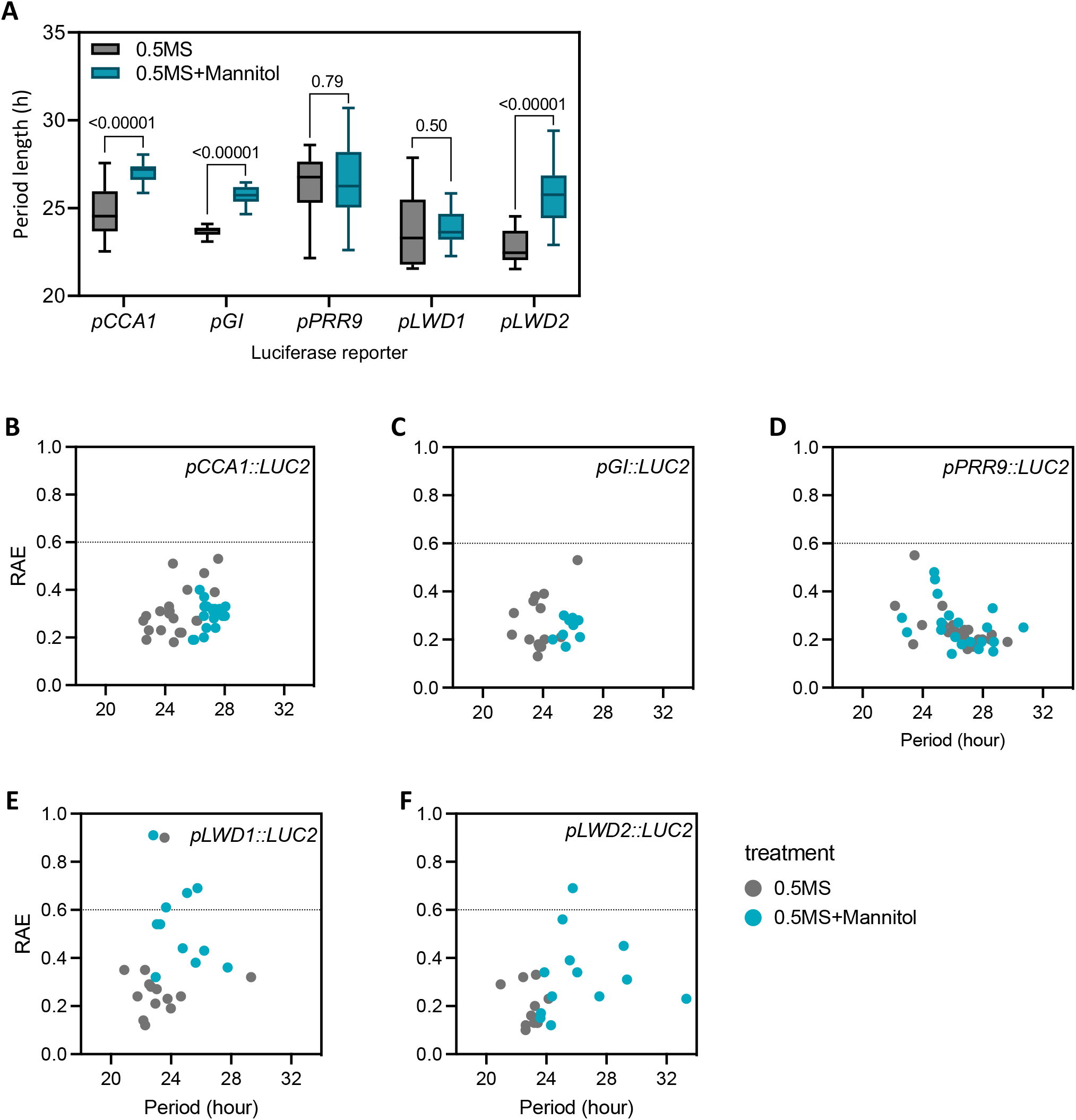
Luciferase reporter constructs highlight varied responses to osmotic stress. **(A)** Circadian free-running period of *pCCA1::LUC2, pGI::LUC2, pPRR9::LUC2, pLWD1::LUC2, pLWD2::LUC2* reporter constructs in the presence or absence of osmotic stress. Plants were grown on 0.5 MS media in 12:12 light:dark cycles for 5 days before transfer to 0.5MS in the presence or absence of 200mM mannitol 24 hours prior to imaging in constant red and blue light (30 μmol m^-2^ s^-1^ and 20 μmol m s respectively). A Mann-Whitney Multiple t-test was used to assess differences in circadian period between treatments. (**B-E**) Assessment of rhythmic robustness (Relative Amplitude Error, RAE) against circadian free-running period for data presented in (A). An RAE of 0 is indicative of a perfect fit whereas an RAE of 1 represents the mathematical limits of rhythm detection (Plautz *et al.*, 1997).

### The SAL1/PAP pathway delays circadian free-running period via cytosolic XRN4

3’ PhosphoAdenosine 5’-Phosphate (PAP) is a retrograde signal that accumulates in response to osmotic stress. Endogenous PAP levels increase as oxidative stress impairs the activity of *SAL1,* a redox-sensitive phosphatase that would otherwise degrade PAP in the chloroplast and mitochondria (Chan *et al.* 2016a, 2016b). *sal1* seedlings have an extended circadian period (Litthauer et al., 2018), and so we were interested whether the accumulation of PAP during osmotic stress varied over the course of a day. PAP accumulation is modest in mock-treated plants, with little variation in PAP levels in these seedlings during the day (Figure 2A, p=0.268). However there was a significant increase in PAP accumulation in mannitol-treated wild-type plants compared to a mock-treated control (p<0.001). By contrast no significant difference in PAP accumulation was observed in plants carrying the *fry1-6* allele of *SAL1* that constitutively accumulate PAP following osmotic stress even though PAP accumulation was substantially higher than that observed in wild-type seedlings [Figure 2A, p=0.24; (Litthauer *et al.*, 2018)]. These data suggest that PAP accumulation during osmotic stress does not vary over the course of the day.

**Figure 2.**
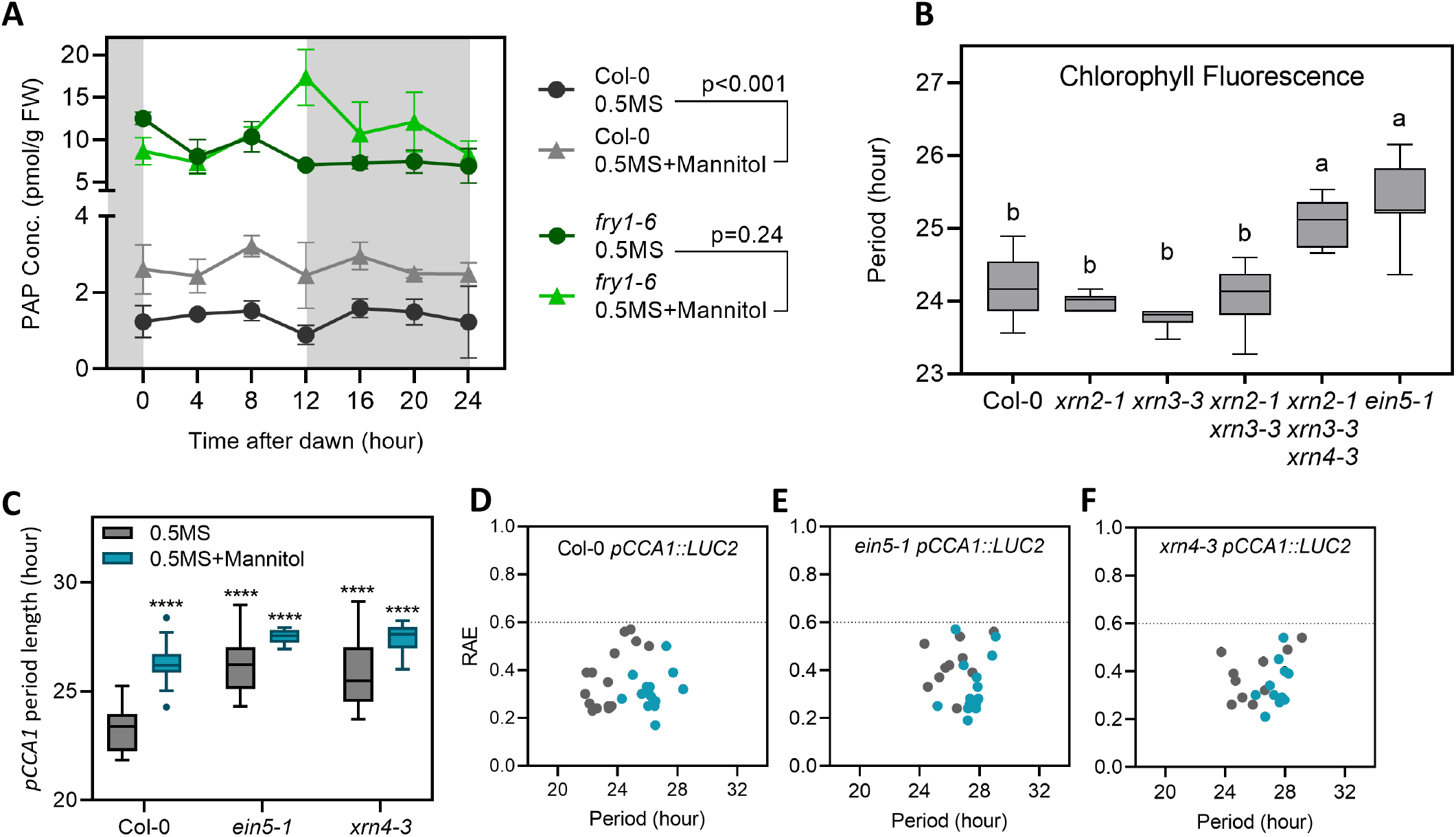
XRN4 contributes towards the extension of circadian free-running period in response to osmotic stress. (**A**) Accumulation of 3’-PhosphoAdenosine 5’-Phosphate (PAP) in Col-0 and *fry1-6* seedlings. Plants were grown on 0.5MS media for 12 d prior to transfer to 200mM mannitol. Seedlings were maintained in entraining conditions prior to harvest on the third day after application of osmotic stress (48-72 hours after transfer). Data are the mean of three biological replicates and analysed using paired t-test; s. e. m. is shown. (**B**) Circadian free-running period of Col-0 (wildtype), *xrn2-1, xrn3-3, xrn2-1xrn3-3, xrn2-1xrn3-3xrn4-3*, and *ein5-1* seedlings were assessed using chlorophyll fluorescence. Seedlings were grown as described in (A) prior to imaging in the absence of osmotic stress. Data were analysed using one-way ANOVA and Tukey’s multiple comparisons test, s. e. m. is shown. (**C**) Circadian free-running period of *xrn4* alleles expressing a *pCCA1::LUC2* reporter construct. Seedlings were grown on 0.5 MS media in 12:12 light:dark cycles for 5 days before transfer to 0.5MS in the presence (blue) or absence (grey) of 200mM mannitol. Data were analysed using one-way ANOVA and Dunnett’s multiple comparisons test (**D-F**) Assessment of rhythmic robustness (Relative Amplitude Error, RAE) against circadian free-running period for data presented in (C), with a threshold set to 0.6.

One of the biochemical consequences of PAP accumulation is the inhibition of the XRN family of exoribonucleases, with Arabidopsis expressing three *XRN* orthologues (Nagarajan *et al.* 2013). *xrn234* seedlings have previously been reported to have extended circadian FRP, although *xrn234* seedlings have a pleiotropic phenotype including impaired growth (Hirsch et al., 2011; Litthauer et al, 2018). We were therefore interested whether specific XRN proteins were sufficient to link the SAL1/PAP signalling pathway into the circadian system. Neither *xrn2-1, xrn3-3,* nor *xrn2-1 xrn3-3* seedlings have a significant extension of free-running period when assessed by chlorophyll fluorescence (Figure 2B, Supplemental Figure 2A). However, we were able to observe that the *ein5-1* allele of *xrn4* did have an impaired circadian system, with a circadian FRP 1 hr longer than wild-type controls that was indistinguishable from *xrn234* seedlings (Figure 2B, Supplemental Figure 2A). A similar extension of FRP was observed using a *pCCA1::LUC2* reporter construct, with both *ein5-1* and *xrn4-3* alleles of *xrn4* having an extended FRP compared to wild type (Figure 2C and Supplemental Figure 2B-G, τ=23.19±0.34hrs, 26.34±0.68hrs, and 26.70±0.60hrs in wild type, *ein5-1* and *xrn4-3* lines respectively). We next assessed whether the circadian system of *xrn4* seedlings retained a response to osmotic stress. *ein5-1* and *xrn4-3* seedlings continued to demonstrate a modest response to osmotic stress, although this was much less pronounced than in wild-type seedlings (Figures 2C-F, Supplemental Figure 2D-G). These data suggest that XRN4 contributes to the extension of circadian FRP in response to osmotic stress, whilst also indicating that additional mechanisms contribute to the integration of osmotic stress into the circadian system.

### XRN4 contributes to the degradation of *LWD1, LWD2,* and *PRR7* transcripts

XRN4 is a cytosolic protein that has roles in co-translational decay and cytosolic 5’-3’ RNA decay (Figure 3A; Kastenmayer and Green, 2000; Nagarajan et al., 2019; Carpentier et al., 2020). Approximately 2000 Arabidopsis transcripts have been proposed as XRN4 substrates following Parallel Analysis of RNA Ends (PARE; Nagarajan et al., 2019). Of these candidate substrates, only three (*PRR7, LWD1,* and *LWD2*) are established components of the circadian system (Figure 3B; Farré EM et al., 2005; Wu et al., 2008). To confirm these RNAseq data, we performed RNA stability assays in the presence of cordycepin to inhibit transcription (Figure 3C-H). *LWD1* was degraded in wild-type seedlings (p < 0.01, Figure 3C), although degradation was not apparent in *ein5-1* (Figure 3D). We were interested to note that *LWD1* RNA degradation was not apparent in either wild type or *ein5-1* plants following the application of osmotic stress (Figures 3C-3D). *LWD2* RNA was also degraded in wild-type seedlings (p < 0.01, Figure 3E), with the transcript being stabilised by osmotic stress (Figure 3E). In contrast to *LWD1, LWD2* transcripts continued to decrease over time in *ein5-1* seedlings, although the rate of degradation was less pronounced than that observed in wild type (Figure 3F). In contrast to *LWD1* and *LWD2, PRR7* was not degraded in wild-type mock-treated samples (Figure 3G). However, we did observe an increase in *PRR7* total RNA relative to control transcripts following osmotic stress (p = 0.01; Figure 3G). These relative increases in *PRR7* transcript may represent residual transcription (although *PRR7* accumulation was greatly reduced compared to a non-cordycepin treated control; Supplemental Figure 3) or indicate that the *PRR7* transcript is more stable than housekeeping genes in these specific conditions. A comparable relative increase in *PRR7* transcript was also observed in mock-treated *ein5-1* seedlings (p = 0.03; Figure 3H), although this increase was not observed in *ein5-1* plants subjected to osmotic stress (Figure 3H). Our data suggest that XRN4 contributes to the degradation of *LWD1, LWD2,* and *PRR7,* although the interaction between XRN4 activity and osmotic stress upon RNA degradation remains to be explored and is likely regulated by additional factors.

**Figure 3.**
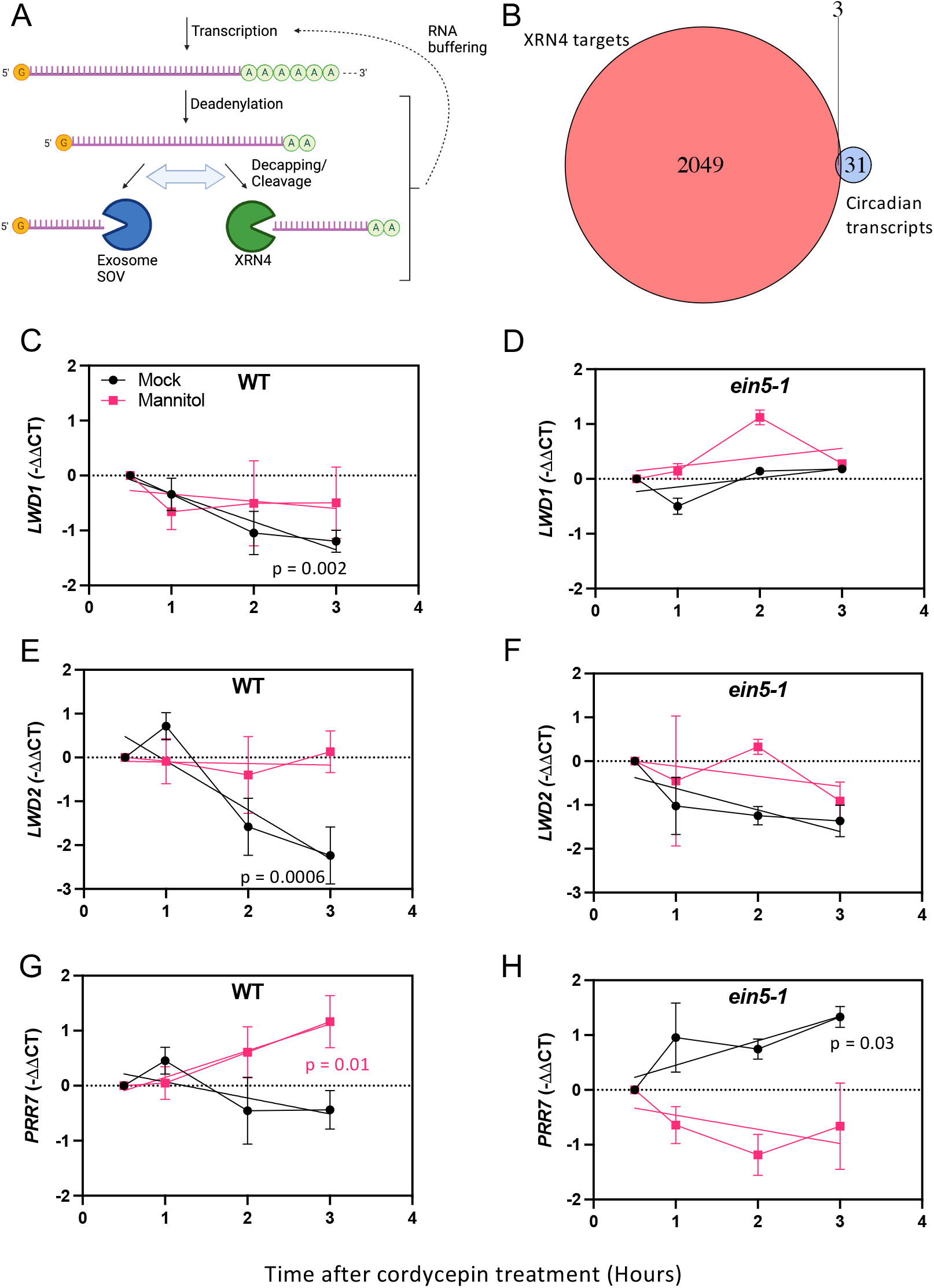
mRNA stability is modulated by osmotic stress and XRN4 activity. **(A)** Outline of RNA degradation pathways in Arabidopsis. Following deadenylation, RNAs are degraded in a 3’-5’ direction by the exosome or SOV (note that Col-0 is an *sov* mutant; Zhang *et al.* 2010). XRN4 degrades RNA in a 5’-3’ direction following endonucleic cleavage or 5’ decapping. Compensatory adjustments between these parallel pathways occur following mutation of ribonucleases, and changes in transcription rates in these cases (‘RNA buffering’) have also been reported (Sorenson *et al.* 2018). Created with BioRender.com. **(B)** Comparison of putative XRN4 degradation targets and characterized components of the circadian system. Putative XRN4 targets were previously identified by Parallel Analysis of RNA Ends (PARE; Nagarajan *et al.* 2019), while defined circadian components were drawn from a previous review (Hsu and Harmer 2014). (**C-D**) Assessment of *LWD1* total transcript stability in Col-0 (C) and *ein5-1* (D) seedlings in the presence or absence of osmotic stress. (**E-F**) Assessment of *LWD2* total transcript stability in Col-0 (E) and *ein5-1* (F) seedlings in the presence or absence of osmotic stress. (**G-H**) Assessment of *PRR7* total transcript stability in Col-0 (G) and *ein5-1* (H) seedlings in the presence or absence of osmotic stress. Seedlings were grown on 0.5MS for 6 days prior to transfer to 200 mM mannitol (pink) or a mock control (black) media. Sampling and application of 0.5 mM cordycepin was completed at ZT4 on day 7; 28 hours after application of osmotic stress. Data are reported relative to transcript accumulation at t=0.5 (30 min after the application of cordycepin). A simple linear regression applied for each combination of genotype and treatment was applied from t=0.5; p-values are shown when the slope is significantly different from 0. Data are the mean of at least three independent experiments, n>10. Error bars indicate s. e. m.

### Osmotic stress disrupts RNA metabolism of specific circadian transcripts

Both transcription and RNA degradation contribute to RNA abundance within cells, with emerging evidence that disruption of cytosolic RNA degradation can influence transcription via ‘RNA buffering’ [Figure 3A; (Sorenson et al., 2018; Sieburth and Vincent, 2018)]. Since RNA degradation is a multi-stage process initiated by deadenylation [Figure 3A; (Sieburth and Vincent, 2018)] we used either oligo-dT (to assess polyadenylated mRNA) or a random hexamer (to capture RNA decay intermediates in addition to polyadenylated mRNAs) following the application of osmotic stress (Figure 4, Supplemental Figure 4). We first examined polyadenylated transcript levels and were interested to note that *CCA1* and *GI* polyadenylated mRNA accumulation remained robust following application of 200 mM mannitol (Figure 4A, Supplemental Figure 4). This contrasts with the general reduction of luciferase bioluminescence observed following osmotic stress (Supplemental Figure 1). We also found that steady state levels of *LWD1* or *LWD2* polyadenylated mRNA were extremely low in either the presence or absence of mannitol despite *pLWD1::LUC* and *pLWD2::LUC* displaying circadian rhythms [Figures 1E-F, 4B-C, (Wang et al., 2011)]. Polyadenylated *PRR7* transcript peak levels were comparable between mock and osmotically stressed seedlings (Figure 4D).

**Figure 4.**
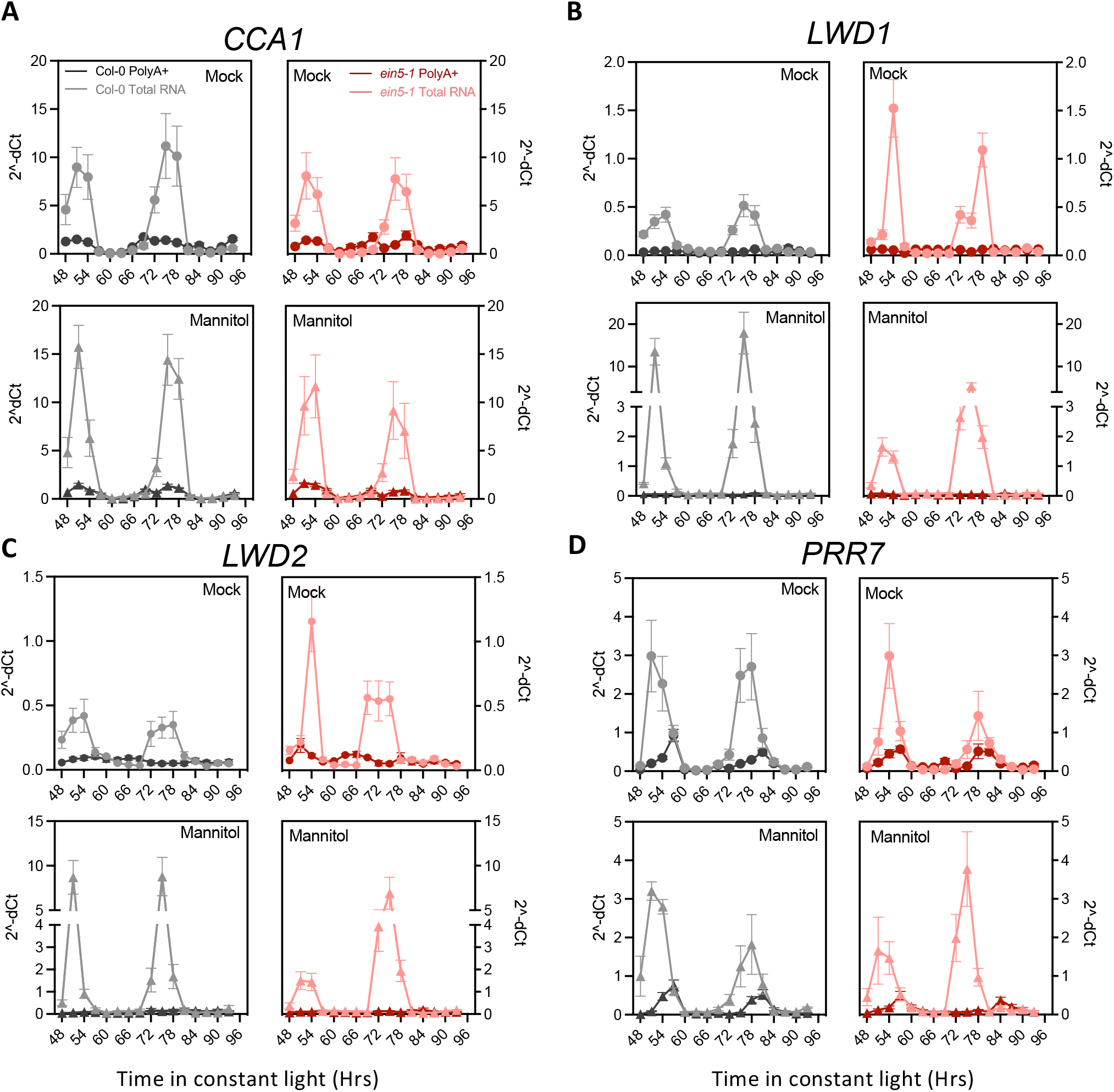
Relative polyadenylated and total RNA abundance of selected circadian clock genes following application of osmotic stress. Wild-type (gray) and *ein5-1* (red) seedlings were grown on 0.5 MS media in 12:12 light:dark cycles for 5 days before transfer to 0.5MS in the presence or absence of 200mM mannitol. Seedlings were returned to entraining conditions for 24 hours prior to transfer to continuous white light (60 μmol m^-2^ s^-1^). Fold-change in *CCA1* (A), *LWD1* (B), *LWD2* (C), and *PRR7* (D) is presented relative to three circadian reference genes listed in Supplemental Table 1. cDNA was synthesized using either an Oligo dT primer or random hexamer to obtain PolyA+ (darker colour) and total transcript (lighter colour), respectively. Data were normalised using 2^-dC_t_ method. Data are representative of at least three independent experiments (n > 10). Error bars indicate s. e. m.

We next compared the accumulation of polyadenylated RNA and total RNA (Figure 4, Supplemental Figure 4). Interestingly, we observed that patterns of polyadenylated RNA did not always correlate with total RNA within cells. For instance, steady-state abundance of *PRR7* and *GI* polyadenylated transcripts peak during the subjective afternoon, whereas total *PRR7* and *GI* RNA begins to accumulate approximately 6 hours earlier (Figure 4D, Supplemental Figure 4). These differences in accumulation patterns were more pronounced for *LWD1* and *LWD2* transcripts, with total RNA of *LWD1* and *LWD2* demonstrating circadian rhythmicity as observed with luciferase reporter lines whereas polyadenylated *LWD1 andLWD2* species did not accumulate (Figures 1E, 1F, 4C-D). The discrepancy between polyadenylated and total *LWD1* and *LWD2* RNA reveals a strong post-transcriptional regulation of these transcripts. We also noted a dramatic increase in total *LWD1* and *LWD2* RNA when wild-type seedlings were subjected to osmotic stress that was not apparent in *ein5-1* (Figures 4B-4C). A dramatic increase in *GI* total RNA was similarly observed following osmotic stress, although this increase was observed in both wild-type and *ein5-1* seedlings (Supplemental Figure 4). By contrast, these increases in total RNA following osmotic stress were not apparent in *PRR7* or *CCA1* species (Figures 4A and 4D), suggesting that *LWD1, LWD2,* and *GI* total RNA species are particularly affected following osmotic stress. These differences between promoter activity, polyadenylated mRNA accumulation, and total RNA (Figures 1 and 4) demonstrate how different aspects of transcript metabolism vary over circadian time.

Since XRN4 contributes to the degradation of a subset of circadian transcripts (Figure 3) we next assessed changes in total RNA accumulation in *xrn4* seedlings. The rhythmic accumulation of *LWD1* and *LWD2* total RNA was retained in *ein5-1* seedlings and was broadly similar to wild type in mock-treated conditions (Figures 4B-4C). Contrary to our hypotheses drawn from RNA stability assays (Figure 3), peak *LWD1* and *LWD2* levels were reduced in *ein5-1* following osmotic stress (compared to the wild type; Figures 4B and 4C). In contrast to *LWD1,* and *LWD2* total RNA, only modest differences in *CCA1*, *PRR7,* and *GI* total RNA were observed in *ein5-1* seedlings compared to wild type controls (Figures 4A, 4D, Supplemental Figure 4). These assessments of total RNA accumulation suggest that circadian accumulation of *LWD1* and *LWD2* total RNAs are greatly influenced by XRN4 activity but do indicate a complex effect beyond simply degrading target transcripts, possibly involving RNA buffering mechanisms (Siebert and Vincent, 2018).

### *PRR7* and *LWD1* enable the circadian response to osmotic stress

We next assessed if *PRR7, LWD1,* or *LWD2* contributed towards tolerance of osmotic stress, or were necessary for the extension of circadian period observed in response to mannitol treatment (Figure 5, Supplemental Figures 5 and 6). All genotypes tested had significantly shorter hypocotyls following the application of 200 mM mannitol (p<0.001; Šídák’s multiple comparisons test), although only *ein5-1* and *xrn4-3* seedlings were significantly shorter than wild type in the presence of osmotic stress (p<0.05, Dunnett’s multiple comparisons test, Supplemental Figure 5). *lwd1 pCCA1::LUC2* seedlings have a short period phenotype in the absence of osmotic stress (Airoldi et al., 2019) but retain the extension of circadian period following the application of mannitol (Figures 5A-5C). *lwd2 pCCA1::LUC2* seedlings demonstrated a longer FRP than wild type during both control and osmotic stress treatments, although the response to osmotic stress remained (Figures 5A and 5D). *lwd1 lwd2 pCCA1::LUC2* seedlings displayed a pronounced shortening of FRP (Wang et al., 2011), but did not display an increase in FRP in response to osmotic stress (p > 0.05, Figures 5A and 5E). We next assessed the contribution of *PRR7* towards the circadian response to osmotic stress. Interestingly, although robust circadian rhythms were maintained in *prr7-3 pGI::LUC2* seedlings transferred to 200 mM mannitol, we did not observe an extension in FRP (Figure 5F-H). These data demonstrate that PRR7 is necessary to maintain the proper response of circadian system to osmotic stress, and suggest that both LWD1 and LWD2 contribute towards the extension of FRP as part of this response.

**Figure 5.**
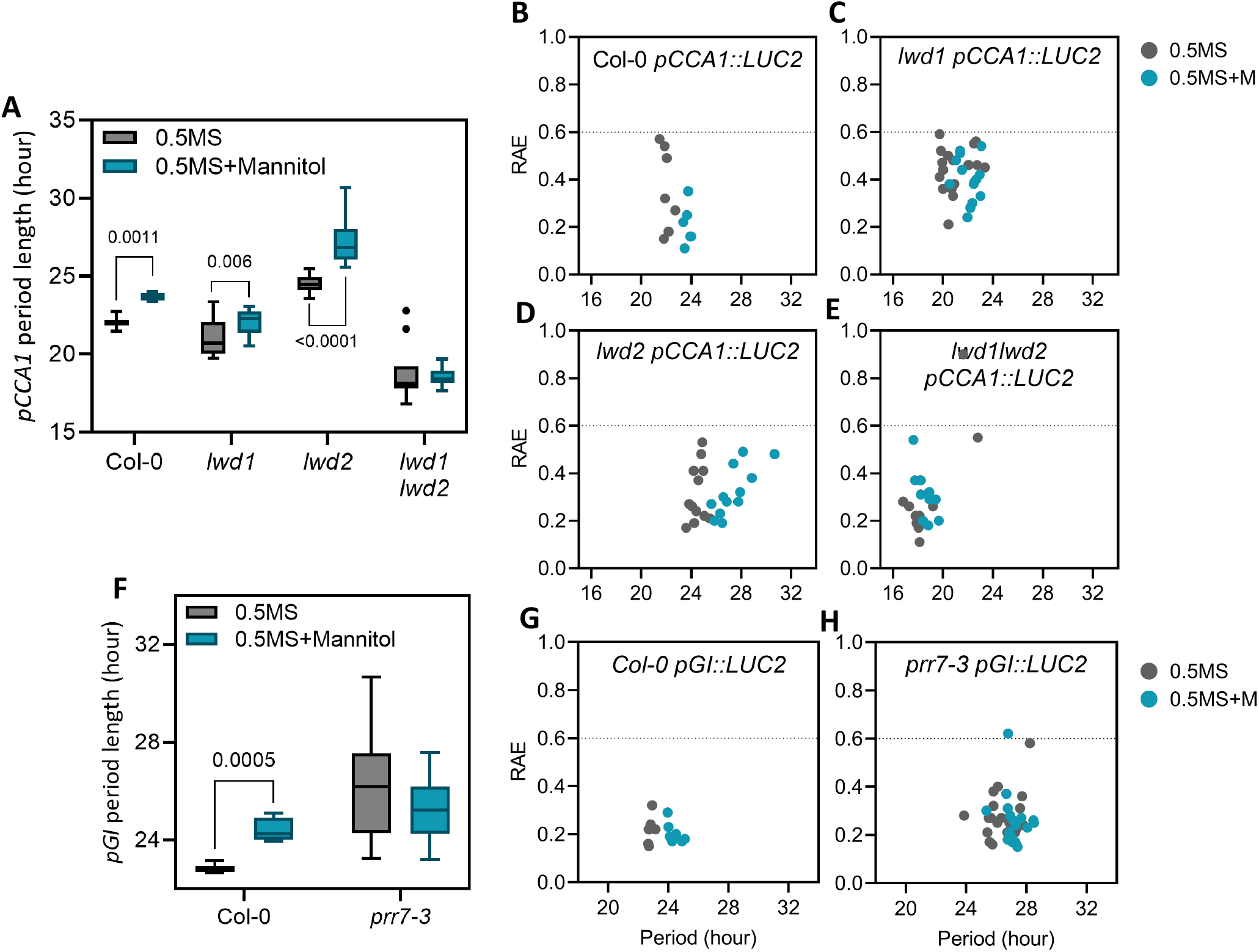
Loss of *LWD1, LWD2,* or *PRR7* perturbs circadian responses to osmotic stress. (**A**) Circadian free-running period of *pCCA1::LUC2* in wildtype, *ein5-1, lwd1*, *lwd2,* and *lwd1lwd2* backgrounds. p-values for the difference in circadian free-running period following osmotic stress is shown (Mann-Whitney Multiple t-test). (**B-E**) Assessment of rhythmic robustness (Relative Amplitude Error, RAE) against circadian free-running period for data presented in (A), with a threshold set to 0.6. (**F**) Circadian free-running period of *pGI::LUC2* reporter lines in wildtype and *prr7-3* backgrounds. p-values for the difference in circadian free-running period following osmotic stress is shown (Mann-Whitney Multiple t-test). (**G,H**) Assessment of rhythmic robustness (Relative Amplitude Error, RAE) against circadian free-running period for data presented in (F), with a threshold set to 0.6. Plants were grown on 0.5 MS media for 5 days and transferred to 0.5MS with or without 200 mM mannitol 24 hours before imaging under constant constant red and blue light (30 μmol m s and 20 μmol m^-2^ s^-1^ respectively).

## Discussion and Conclusions

### Post-transcriptional regulation distinguishes the accumulation of polyadenylated mRNA from circadian patterns of promoter activity and RNA decay intermediates

Our initial experiments using luciferase bioluminescence reporters suggested that individual luciferase reporters within the circadian system were differentially regulated in response to osmotic stress, with *LWD1* and *PRR9* promoter-driven lines lacking a circadian response to the application of osmotic stress (Figure 1). Divergence between different luciferase reporter behavior have previously been reported and may arise in part from tissuespecific expression patterns (Endo *et al.*, 2014, Nimmo et al., 2020, Hall *et al.*, 2002, Haydon *et al.*, 2013). However we were interested whether these differences were reflective of the circadian system adapting to osmotic stress.

Although modern circadian molecular biology is founded upon luciferase reporter constructs that reveal circadian rhythms of reporter activity (Millar et al., 1992), post-transcriptional regulation of some transcripts has been apparent for several years. For example, uniform levels of *LIGHT HARVESTING CHLOROPHYLL BINDING PROTEIN (LHCB1*3*) transcript are maintained despite luciferase activity driven from this promoter being rhythmic (Millar and Kay, 1991; Millar et al., 1992). Conversely, *NITRATE REDUCTASE2* is transcribed constantly, and yet has rhythmic mRNA accumulation (Pilgrim *et al.*, 1993). In our study, it is apparent polyadenylated *LWD1* RNA is arrhythmic in constant light, despite a *pLWD1::LUC* reporter line and *LWD1* total RNA displaying circadian regulation (Figures 1 and 4). A similar phenotype is observed for *LWD2* transcripts (Figures 1 and 4). The length of the polyadenylated tail correlates negatively with gene expression in Arabidopsis (Parker et al., 2020), while removal of the polyadenylation signal is an important initial step in RNA degradation (Figure 3A, Sieburth and Vincent, 2018). The lack of significant accumulation of polyadenylated *LWD1* and *LWD2* in constant light may therefore indicate that *LWD1* and *LWD2* are rapidly transcribed and degraded, although additional experimentation will be required to test this hypothesis. We were also interested to note that peak accumulation of *PRR7* total RNA preceded the phase of peak polyadenylated *PRR7* RNA by several hours (Figure 4D). Differences in post-transcriptional processing presumably contribute to disparities we observe between luciferase reporter activity and steady-state transcript accumulation in response to osmotic stress (Figures 1, 4 and 5, Supplemental Figures 1 and 6).

### RNA degradation via XRN4 contributes to the maintenance of circadian rhythms

3’-PhosphoAdenosine 5’-Phosphate (PAP) accumulates during osmotic stress, leading to the inhibition of exoribonuclease activity [Figure 3A, (Dichtl et al., 1997; Estavillo et al., 2011; Litthauer et al., 2018)]. The accumulation of PAP precipitates an extension of circadian period through the inactivation of XRN exoribonucleases, although it was not apparent from previous studies whether this phenotype was due to increased RNA Polymerase II 3’ read-through resulting from the nuclear-localised XRN2 and XRN3 or instead arose from catabolism in the cytosol via XRN4 (Crisp et al., 2018; Litthauer et al., 2018; Carpentier et al., 2020). Our studies demonstrate that the loss of XRN4 is sufficient to induce extension of circadian FRP (Figure 2C-F), highlighting the contribution of cytosolic RNA degradation to the maintenance of circadian rhythms. XRN4 contributes to the degradation of *PRR7* and *LWD1* although circadian rhythms of *PRR7, LWD1,* and *LWD2* total RNA were still observed in *ein5-1* seedlings [Figures 3 and 4, (Nagarajan et al., 2019)]. While we cannot exclude a role for XRN4 in the degradation of additional circadian transcripts, further timepoints harvested throughout the diel cycle will be necessary to obtain a comprehensive dataset describing all clock genes regulated by XRN4 as *XRN4* accumulation is constant in both long-day and short-day conditions (Litthauer et al., 2018). Indeed, it will also be of interest to understand the contribution of XRN4 towards the translation of circadian proteins during osmotic stress, given the contribution of XRN4 to co-translational decay (Carpentier et al., 2020).

The application of osmotic stress induces changes in both steady-state RNA accumulation and RNA degradation rates (Figures 3 and 4). Although the differences in total RNA accumulation may be mediated in part via the PAP-mediated inhibition of XRN4 activity, it is apparent that XRN4 is not the sole contributor to RNA stability (Figures 3 and 4). For example, the unanticipated reduced accumulation of *LWD1* and *LWD2* total RNA at peak times in *ein5-1* seedlings indicate that additional factors (such as transcriptional regulation or compensatory RNA degradation mechanisms) are also perturbed by the loss of XRN4 activity (Figure 4; Liu and Chen, 2016). In yeast, the XRN4 functional orthologue XRN1 couples transcription with RNA decay, shuttling into the nucleus as part of a feedback mechanism to regulate mRNA accumulation and translation (Blasco-Moreno et al., 2019; Haimovich et al., 2013; Sun et al., 2013). While this latter mechanism has not been explicitly reported in plants, global analysis of RNA decay in *sov* seedlings reveals communication between cytoplasmic decay and the transcriptional machinery (Sorenson et al., 2018; Sieburth and Vincent, 2018). Although the activity of XRN4 only accounts for a portion of the phenotypes observed, these data support the hypothesis that the circadian system adapts to osmotic stress through post-transcriptional regulation. Additional work is also needed to understand which of these changes are directly precipitated by osmotic stress rather than being an indirect consequence of circadian perturbation.

### *PRR7, LWD1,* and *LWD2* contribute to circadian responses to osmotic stress

Although little has been reported regarding the contribution of LWD1 and LWD2 to abiotic stress, PRR7 is increasingly recognized as a crucial circadian component that contributes metabolic information to the molecular timekeeper (Liu et al., 2013, Webb et al., 2019). Since *prr7-3* and *lwd1 lwd2* seedlings demonstrate impaired circadian responses to osmotic stress [Figure 5, (Nagarajan et al., 2019)] it is possible that altered accumulation of total *LWD1,* and *LWD2* transcripts following osmotic stress (Figure 4) contributes to the observed extension of circadian FRP, although the biological function of these RNAs remains to be determined given that changes in polyadenylated RNA remains modest in constant light (Figure 4).

Previous work has reported enrichment of PRR7 targets that are responsive to abiotic stress (Liu et al., 2013). Indeed, the preponderance of reports linking PRR7 to abiotic stress responses suggest that PRR7 is a central component of plants’ responses to abiotic stresses such as heat, shade and drought (Liu et al., 2013; Kolmos et al., 2014; Blair et al., 2019; Zhang et al., 2020). A high percentage (28%) of PRR7 targets are also ABA-regulated, with more than one third of PRR7 target genes possessing ABA-responsive elements (Liu et al., 2013). It is therefore possible that regulation of *PRR7* by XRN4 provides an additional pathway for PAP to modulate ABA-induced signaling (Pornsiriwong et al., 2017). Such a notion would also align with the proposed role of PRR7 as a dynamic integrator of photosynthetic signals into the circadian system (Webb et al., 2019) and underscore the importance of PRR7 as an integrator of environmental signals.

## Materials and Methods

### Plant material, growth, and treatments

Plant genotypes used in this work are listed in Supplemental Table 1. Plants were germinated and grown on half-strength Murashige and Skoog (0.5 MS) media for 5 days before being transferred to either half-strength 0.5 MS media or 0.5 MS supplemented with 200mM mannitol as indicated. Plants were grown under 60 μmol m ^-2^ s^-1^ white light in 12:12 light:dark cycles. Relative humidity and temperature were set to 60%-70% and 22°C, respectively.

### Hypocotyl measurements

Seedlings were germinated on 0.5MS media and grown under 60 μmol m^-2^ s^-1^ white light in 8:16 light:dark cycles for three days prior to transfer to plates containing 200 mM mannitol or a mock-treated control.

Hypocotyl length was measured at 7 days after germination using ImageJ (Abramoff *et al.* 2004).

### Luciferase activity

Seedlings were grown in 12:12 light:dark cycles before being sprayed with 3mM d-luciferin in 0.01% Triton X-100 before being returned to entraining conditions for 24 hours. The age of seedlings used in each experiment is described in the respective figure legend. Luciferase imaging was completed under constant light conditions (20 μmol m^-2^ s^-1^ constant blue and 30 μmol m^-2^ s^-1^ constant red light) for 5 days. Images were taken every two hours with a QImaging Retiga LUMO Monochrome Camera controlled by a MicroManager 1.4 script. Circadian parameters were determined using the website biodare2.ed.ac.uk which used Fourier fast transform-nonlinear least squares to calculate circadian parameters (Moore et al., 2014).

### Chlorophyll fluorescence imaging

Chlorophyll fluorescence parameters were recorded with a Fluorimager imaging system (Technologica) as previously described (Litthauer et al., 2015). Patterns of *F_q_′/F_m_′* were fitted to cosine waves using FFT-NLLS (Plautz et al., 1997) to estimate circadian period length and additional circadian parameters. Sample size was chosen to achieve a power of 0.8 in a two-sample t test at α = 0.05. Previously collected data were used to estimate σ = 0.6.

### Assessment of PAP accumulation

Twelve-day-old seedlings grown on 0.5MS were transferred to 0.5MS medium supplemented with either 200mM mannitol or a mock control. Seedlings were returned to light:dark cycles for two days prior to harvesting on day 3 of osmotic stress at 4h intervals and stored at −80□C until processing. Plant tissue was ground using a TissueLyser (Qiagen-Retsch) and then incubated in 0.1M HCl for 15’. Particulates were precipitated twice by centrifugation and the supernatant was added to CP buffer (620 mM citric acid, 760 mM Na_2_HPO_4_, pH 4). The samples were then derivitised with chloroacetyl-aldehyde at 80□C for 10’ prior to measurement using an HPLC system (Shimadzu) with a Phenomenex Luna 5μm C18(2) 100Å LC 150×4.6mm column. The column was equilibrated with 97%(v/v) Buffer A (5.7mM [CH_3_(CH_2_)_3_]_4_NHSO_4_ and 30.5 mM KH_2_PO_4_, pH 5.8) and 3%(v/v) acetonitrile, after injection the concentration of acetonitrile rose to 33%(v/v) with a linear gradient across 43’20”, the column was then re-equilibrated with 97%(v/v) Buffer A and 3%(v/v) acetonitrile for 6’40”. Concentration of PAP was measured relative to commercially available standards.

### Gene expression analyses

For gene expression, 10-15 seedlings were pooled. RNA isolation was performed using TriZol™ (Sigma) based on the manufacturers’ instructions. DNA contaminant was removed using DNase I - RNase-free (Thermo Scientific™) and cDNA was synthesized using RevertAid First Strand cDNA Synthesis Kit (Thermo Scientific™) with Oligo dT or random hexamer as specified in the figure legend. The resultant cDNA was used as template for real-time PCR (primers listed in Supplemental Table 2) using StepOne™ Real-Time PCR System (Applied Biosystems™). Data were processed using the dC_t_ method, and is presented relative to *APA1, APX3,* and *IPP2* which have previously been reported to be stable over circadian time (Nusinow *et al.* 2011).

### RNA stability assay

RNA stability was assessed as previously described (Sorenson et al., 2018). Seedlings were preincubated in 3 mL incubation buffer (15 mM sucrose, 1mM PIPES pH 6.25, 1mM KCl, 1mM sodium citrate) with aeration (swirling 100 rpm) in petri dishes for 15 min. Transcription was inhibited by adding 3 mL of fresh buffer containing 1 mM cordycepin. Vacuum infiltration was performed for 15 mins. Tissue was collected 30, 60, 120, and 180 min after vacuum release, snap-frozen in liquid nitrogen, and stored at −80 °C prior to RNA extraction. cDNA was synthesized using a random hexamer oligo. RNA steady-state accumulation is presented relative to *GAMMA SUBUNIT OF MT ATP SYNTHASE* (*ATP3*), *EUKARYOTIC ELONGATION FACTOR 5A-2* (*ELF5A-2*), and *PLASTID ISOFORM TRIOSE PHOSPHATE ISOMERASE* (*PDTPI*), which have previously been reported to be stable during RNA degradation assays (Sorenson *et al.* 2018).

## Accession numbers

Genes examined in this article can be found in the Arabidopsis Genome Initiative database under the following accession numbers: *APA1,* At1g11910; *APX3,* At4g35000; *ATP3,* At2g33040; *CCA1*, At2g46830; *ELF4,* At2g40080; *ELF5A-2,* At1g26630*; GIGANTEA,* At1g22770; *IPP2*, At3g02780; *LHY*, At1g01060; *LWD1,* At1g12910; *LWD2,* At3g26640; *PDTPI,* AT2g21170; *PRR7,* At5g02810; *PRR9,* At2g46790; *SAL1,* At5g63980; *TOC1,* At5g61380;*XRN2,* At5g42540;*XRN3,* At1g75660;*XRN4,* At1g54490.

## Acknowledgements

The authors thank Dr James Locke (Sainsbury Laboratory, University of Cambridge, UK), Prof. Alex Webb (University of Cambridge, UK), and Prof. Wu (Academia Sinica, Taiwan) for the provision of seed.

---

**Supplemental Figure 1. Raw and Normalised bioluminescence waveforms of data presented in Figure 1;** (**A-B**)*pCCA1::LUC2,* (**C-D**)*pGI::LUC2*, (**E-F**)*pPRR9::LUC2,* (**G-H**)*pLWD1::LUC2,* (**I-J**)*pLWD2::LUC2.* Data were normalized using BioDare2 (Moore *et* al. 2014). Mean ± s. e. m. are shown.

**Supplemental Figure 2. Assessment of circadian rhythms in *xrn* seedlings. (A)** Circadian rhythms of chlorophyll fluorescence under constant light. Period estimates calculated from these data are shown in Figure 2B. **(B-G)** Raw (B,D,F) and normalized (C, E, G) bioluminescence waveforms data of *pCCA1::LUC2* presented in Figure 2C-F. (B-C) Col-0 (wildtype), (D-E) *ein5-1,* and (F-G) *xrn4-3* background. Data were normalized using BioDare2 (Moore *et* al. 2014). Mean ± s. e. m. are shown. Waveforms presented are representative of three independent experiments.

**Supplemental Figure 3.** Assessment of *PRR7* transcript accumulation in wild-type and *ein5-1* seedlings in the absence of cordycepin to inhibit transcription. Data are the mean of n>3. Error bars indicate s. e. m.

**Supplemental Figure 4. Relative polyadenylated and total RNA abundance of *GIGANTEA* following application of osmotic stress.** Wild-type (gray) and *ein5-1* (red) seedlings were grown on 0.5 MS media in 12:12 light:dark cycles for 5 days before transfer to 0.5MS in the presence or absence of 200mM mannitol. Seedlings were returned to entraining conditions for 24 hours prior to transfer to continuous white light (60 μmol m^-2^ s^-1^). Fold-change in *GIGANTEA* is presented relative to three circadian reference genes listed in Supplemental Table 1. cDNA was synthesized using either an Oligo dT primer or random hexamer to obtain PolyA÷ (darker colour) and total transcript (lighter colour), respectively. Data were normalised using 2^-dC_t_ method. Data are representative of at least three independent experiments (n > 10). Error bars indicate s. e. m.

**Supplemental Figure 5. Hypocotyl lengths of seedlings in the presence or absence of osmotic stress.** Seedlings were germinated on 0.5x MS media for three days prior to transfer to either 200 mM mannitol or a mock control. Seedlings were measured three days after transfer (six days after germination). All genotypes tested had significantly shorter hypocotyls following the application of 200 mM mannitol (p<0.001; Šídák’s multiple comparisons test). Asterisks indicate significant differences from wild type seedlings in either mock (black asterisks) or stressed conditions (pink asterisks) using Dunnett’s multiple comparisons test (p<0.05).

**Supplemental Figure 6. Normalised bioluminescence waveforms of data presented in Figure 5.** (**A-D**) *pCCA1::LUC2* reporter in (A) Col-0 (wildtype), (B) *lwd1,* (C) *lwd2,* and (D) *lwd1lwd2* background. (**E,F**) *pGI::LUC2* reporter in (**E**) Col-0 (wildtype) and (**F**) *prr7-3* background. Data were normalized using BioDare2 (Moore *et* al. 2014). Mean ± s. e. m. are shown. Waveform data are representative of three independent experiments.

**Supplemental Table 1.** Plant genotypes used in this work.

**Supplemental Table 2.** Oligos used for qRT-PCR.

