## Supplemental for "EXORIBONUCLEASE4 integrates metabolic signals induced by osmotic stress into the circadian system"

### Supplemental Figure 1

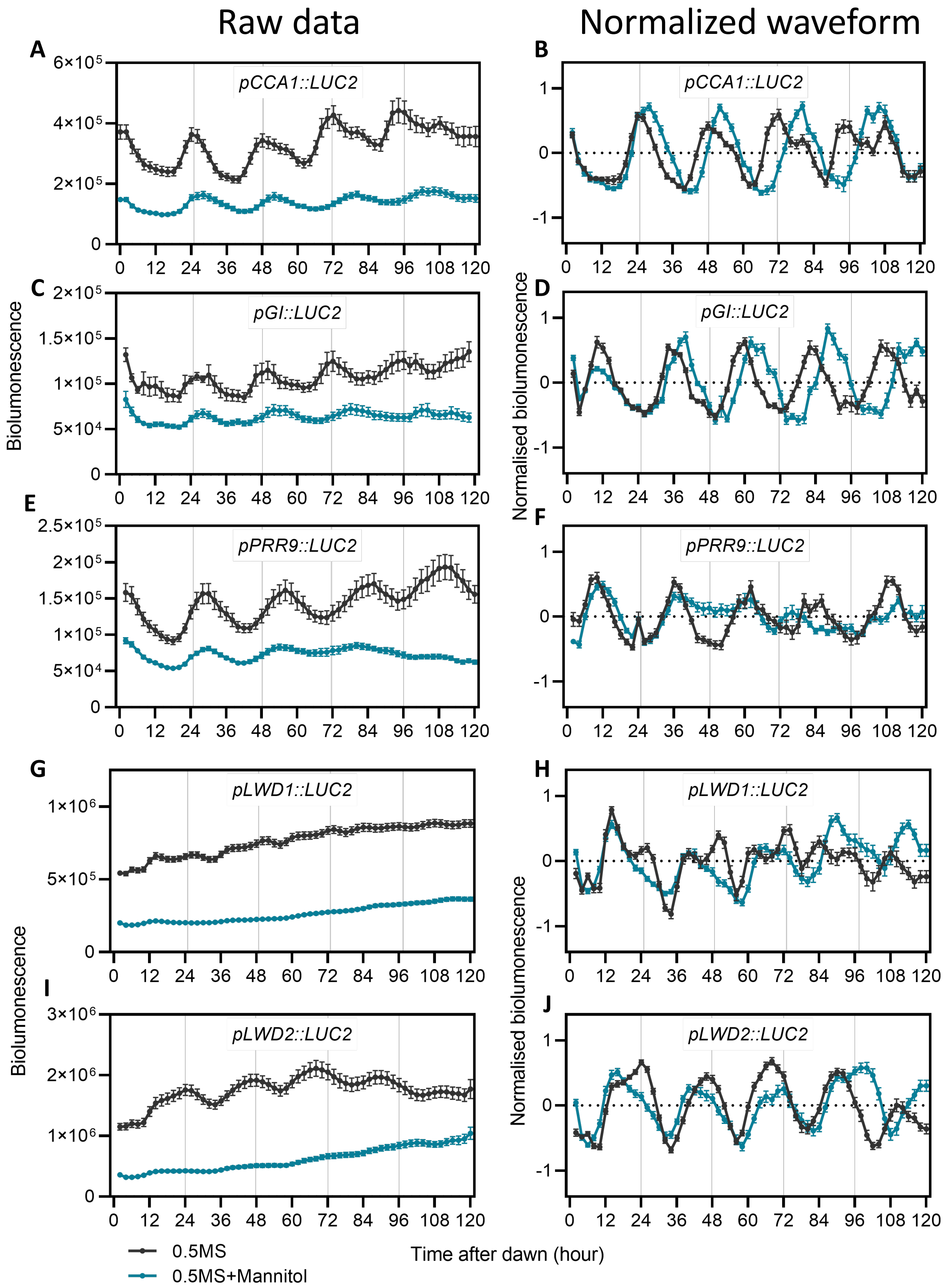


**Supplemental Figure 1. Raw and Normalised bioluminescence waveforms of data presented in Figure 1;** (**A-B**) *pCCA1::LUC2*, (**C-D**) *pGI::LUC2* , (**E-F**) *pPRR9::LUC2*, (**G-H**) *pLWD1::LUC2*, (**I-J**) *pLWD2::LUC2*. Data were normalized using BioDare2 (Moore *et* al. 2014). Mean ± s. e. m. are shown.

### Supplemental Figure 2

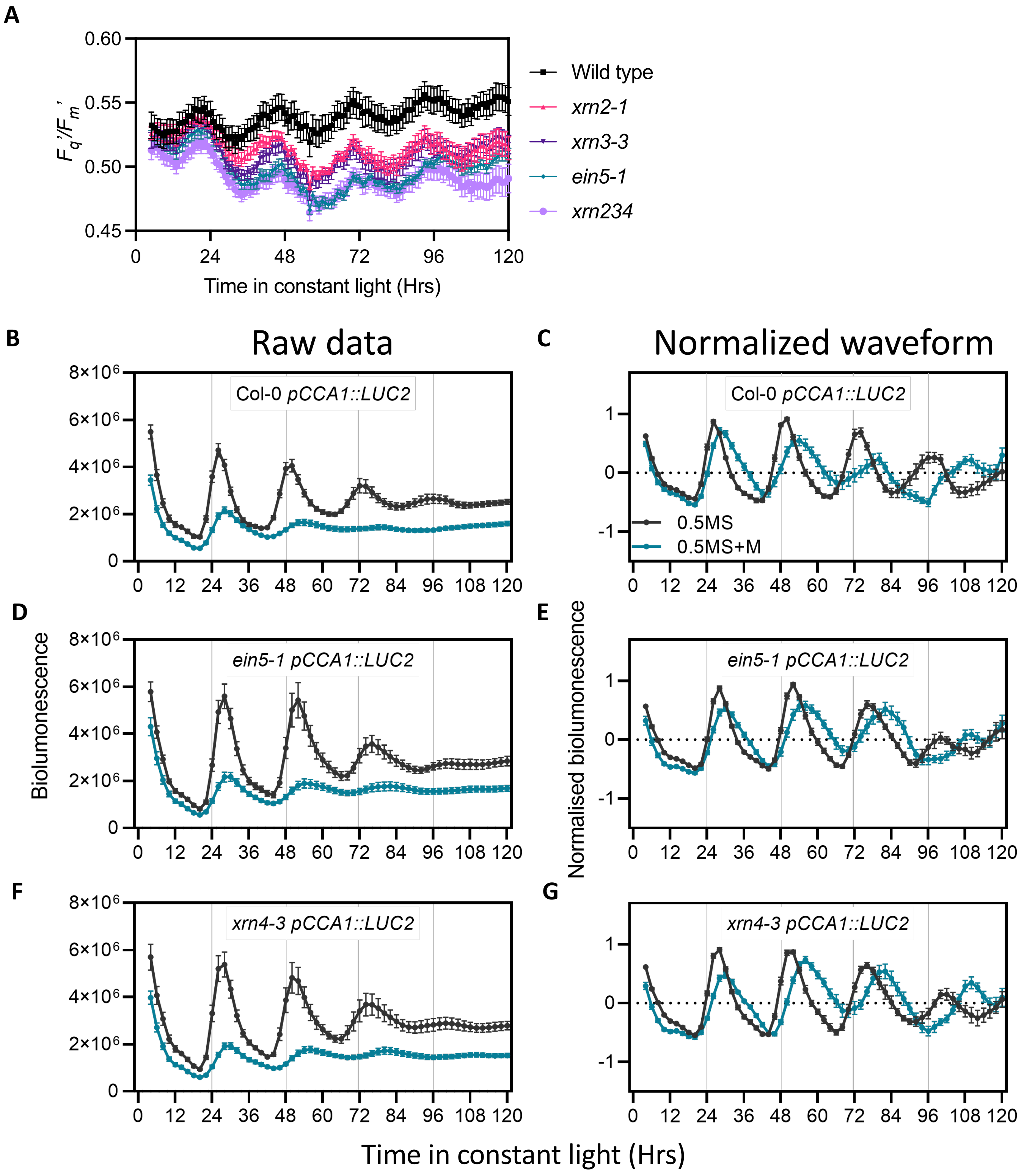


**Supplemental Figure 2. Assessment of circadian rhythms in *xrn* seedlings. (A)** Circadian rhythms of chlorophyll fluorescence under constant light. Period estimates calculated from these data are shown in Figure 2B. **(B-G)** Raw (B,D,F) and normalized (C, E, G) bioluminescence waveforms data of *pCCA1::LUC2* presented in Figure 2C-F. (B-C) Col-0 (wildtype), (D-E) *ein5-1*, and (F-G) *xrn4-3* background. Data were normalized using BioDare2 (Moore *et* al. 2014). Mean ± s. e. m. are shown. Waveforms presented are representative of three independent experiments.

### Supplemental Figure 3

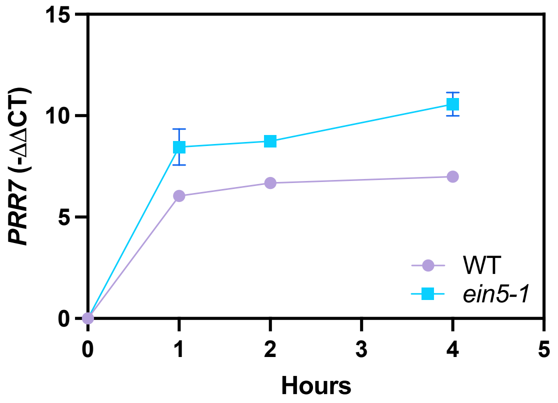


**Supplemental Figure 3.** Assessment of *PRR7* transcript accumulation in wild-type and *ein5-1* seedlings in the absence of cordycepin to inhibit transcription. Data are the mean of n>3. Error bars indicate s. e. m.

### Supplemental Figure 4

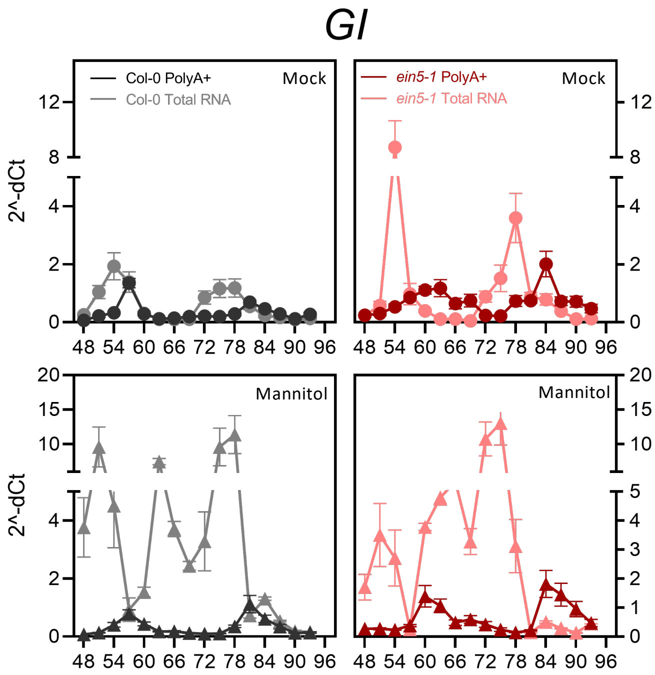


**Supplemental Figure 4. Relative polyadenylated and total RNA abundance of *GIGANTEA* following application of osmotic stress.** Wild-type (gray) and *ein5-1* (red) seedlings were grown on 0.5 MS media in 12:12 light:dark cycles for 5 days before transfer to 0.5MS in the presence or absence of 200mM mannitol. Seedlings were returned to entraining conditions for 24 hours prior to transfer to continuous white light (60 µmol $m^{-2}$ $s^{-1}$). Fold-change in *GIGANTEA* is presented relative to three circadian reference genes listed in Supplemental Table 1. cDNA was synthesized using either an Oligo dT primer or random hexamer to obtain PolyA+ (darker colour) and total transcript (lighter colour), respectively. Data were normalised using 2^-d$C_t$ method. Data are representative of at least three independent experiments (n > 10). Error bars indicate s. e. m.

### Supplemental Figure 5

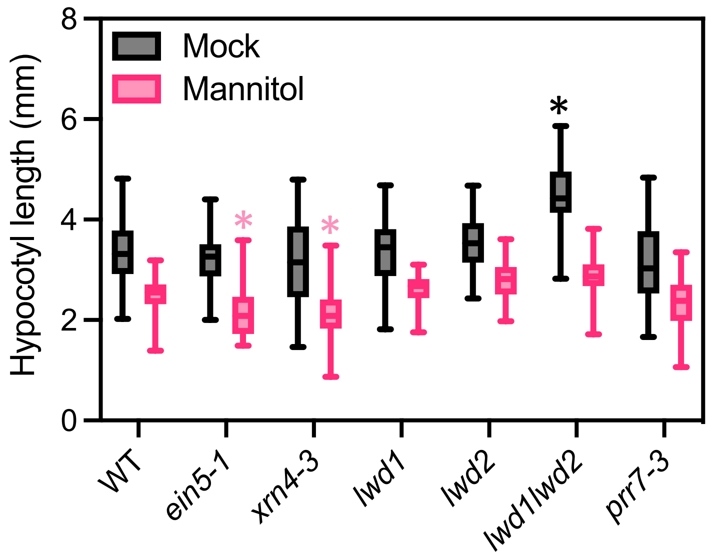


**Supplemental Figure 5. Hypocotyl lengths of seedlings in the presence or absence of osmotic stress.** Seedlings were germinated on 0.5x MS media for three days prior to transfer to either 200 mM mannitol or a mock control. Seedlings were measured three days after transfer (six days after germination). All genotypes tested had significantly shorter hypocotyls following the application of 200 mM mannitol ($p<0.001$; Šídák's multiple comparisons test). Asterisks indicate significant differences from wild type seedlings in either mock (black asterisks) or stressed conditions (pink asterisks) using Dunnett’s multiple comparisons test ($p<0.05$).

### Supplemental Figure 6

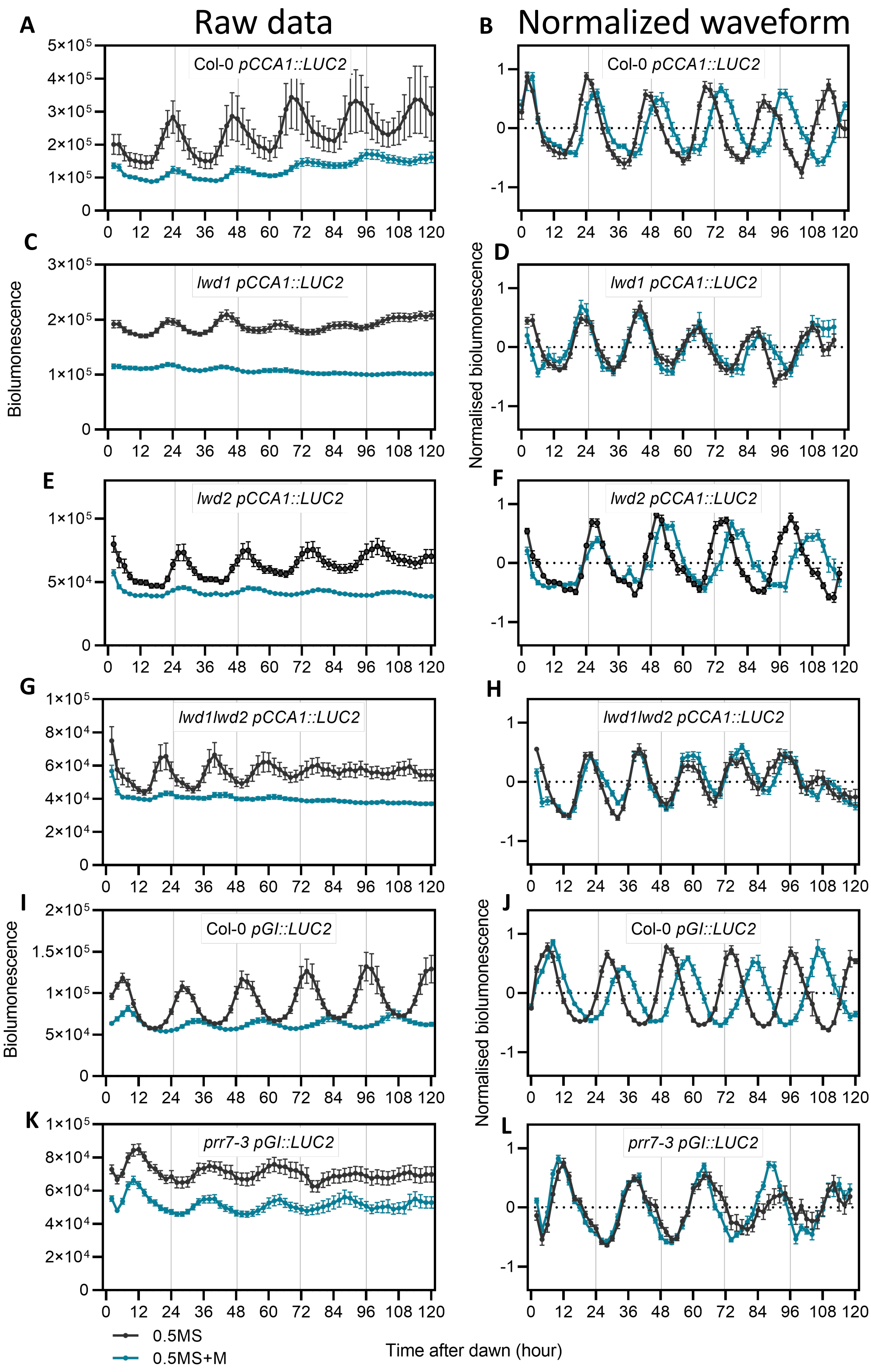


**Supplemental Figure 6. Raw and Normalised bioluminescence waveforms of data presented in Figure 5.** (**A-D**) *pCCA1::LUC2* reporter in (A-B) Col-0 (wildtype), (C-D) *lwd1*, (E-F) *lwd2*, and (G-H) *lwd1lwd2* background. (**I-L**) *pGI::LUC2* reporter in (**I-J**) Col-0 (wildtype) and (**K-L**) *prr7-3* background. Data were normalized using BioDare2 (Moore *et* al. 2014). Mean ± s. e. m. are shown. Waveform data are representative of three independent experiments.

### Supplemental Table 1

| Genotype | Target gene/reporter | Description |
|---|---|---|
| *xrn2-1* | *XRN2* | Gy *et al.* 2007 |
| *xrn3-3* | *XRN3* | Gy *et al.*2007 |
| *fry1-6* | *FRY1/SAL1* | SALK_020882, Litthauer *et al.* 2018 |
| *Col-0* | *GI::LUC* | Locke *et al*. 2006 |
| *prr7-3* | *GI::LUC* | Greenwood *et al.* 2019 |
| *Col-0* | *CCA1::LUC2* | Jones *et al*. 2015 |
| *xrn4-3* | *CCA1::LUC* | SALK_014209, Potuschak *et al.* 2006, this study |
| *ein5-1* | *CCA1::LUC* | Olmedo *et al*. 2006, this study |
| *lwd1* | *CCA1::LUC* | Airoldi *et al.* 2019 |
| *lwd2* | *CCA1::LUC* | Airoldi *et al.* 2019 |
| *lwd1lwd2* | *CCA1::LUC* | Airoldi *et al.* 2019 |
| *Col-0* | *PRR9::LUC* | Locke *et al*. 2006 |
| *Col-0* | *LWD1::LUC* | Wu *et al.* 2008 |
| *Col-0* | *LWD2::LUC* | Wu *et al.* 2008 |

**Supplemental Table 1.** Plant genotypes used in this work

### Supplemental Table 2

| GENE | AGI | Experiment | Forward 5'-3' | Reverse 3'-5' |
|---|---|---|---|---|
| *TOC1* | AT5G61380 | Circadian timecourse | AATAGTAATCCAGCGCAATTTTCTTC | CTTCAATCTACTTTTCTTCGGTGCT |
| *CCA1* | AT2G46830 | Circadian timecourse | CAGCTCCAATATAACCGATCCAT | CAATTCGACCCTCGTCAGACA |
| *PRR7* | AT5G02810 | Circadian timecourse and RNA degradation | GAATGTGCTGAGGCGTTCAGA | GGCTGGATTATACCTTGAGAAAGC |
| *LWD1* | AT1G12910 | Circadian timecourse and RNA degradation | GACCCTATTCTAGCTTACACTGC | ACCCTGAGAATTTGCAGCTTAGT |
| *LWD2* | AT3G26640 | Circadian timecourse and RNA degradation | CCATTAGAAGAAGAAGACGAAGC | CATTTTGGATTTGATCGCTGCT |
| *APX3* | AT4G35000 | Reference for circadian timecourses | GCCGTGAGCTCCGTTCTCT | TCGTGCCATGCCAATCG |
| *IPP2* | AT3G02780 | Reference for circadian timecourses | GTATGAGTTGCTTCTGGAGCAAAG | GAGGATGGCTGCAACAAGTGT |
| *APA1* | AT1G11910 | Reference for circadian timecourses | CTCCAGAAGAGTATGTTCTGAAAG | TCCCAAGATCCAGAGAGGTC |
| *ATP3* | At2g33040 | Reference for RNA degradation assays | GAGGGTGAGACGGTCGAG | GTTCCAATTCTCTTGTGTGATGTTC |
| *ELF5A-2* | AT1G26630 | Reference for RNA degradation assays | ATGGCTTCGTGAGCCTTCTC | CATGACAGACACCACAATATCCTTTC |
| *PDTPI* | AT2G21170 | Reference for RNA degradation assays | CTGTTCCACGCTGATTTCAC | GTTGTTGTGTCACCTCCATTTG |

**Supplemental Table 2.** Oligos used for qRT-PCR.
